# Molecular Transport of the Zika Virus by the Human Cytoplasmic Dynein-1

**DOI:** 10.1101/2022.06.15.496315

**Authors:** Dan Israel Zavala Vargas, Giovani Visoso Carbajal, Leticia Cedillo Barrón, Jessica Georgina Filisola Villaseñor, Romel Rosales Ramirez, Juan E. Ludert, Edgar Morales-Ríos

**Affiliations:** Department of Biochemistry, Center for Research and Advanced Studies (Cinvestav), Mexico City 07360, Mexico; Department of Molecular Biomedicine, Center for Research and Advanced Studies (Cinvestav), Mexico City 07360, Mexico; Department of Infectomics and Molecular Pathogenesis, Center for Research and Advanced Studies (Cinvestav), Mexico City 07360, Mexico

**Keywords:** Zika virus, ZIKV, dynein, envelope protein

## Abstract

Zika virus (ZIKV) infection is a major public health threat, making the study of its biology a matter of great importance. By analyzing the viral-host protein interactions and proposing them as new drug targets, we would diminish the emergence of new resistant strains. In this work, we have shown that human cytoplasmic dynein-1 (Dyn) interacts with the ZIKV. We additionally demonstrate that the envelope protein of the ZIKV and the dimerization domain of the heavy chain of Dyn binds directly without dynactin or cargo adaptor. In addition, we have analyzed this interaction in Vero cells, where we are proposing that the interaction ZIKV-Dyn is finely tuned within the replication cycle. Altogether, our data suggest a new step in the previously described replication cycle of the ZIKV, introducing a suitable molecular target to modulate infection by ZIKV.

## Introduction

Several flaviviruses are neurotropic (for example, WNV, JEV, TBEV, USUV, ZIKV and ILHV), it can spread to the brain and spinal cord and cause severe neurological syndromes including meningitis, encephalitis and acute flaccid paralysis. Infections caused by these viruses can result in death or long-term disability in survivors. Other flaviviruses (such as YFV, DENV and ZIKV) cause visceral disease resulting in liver failure, hemorrhagic syndromes and vascular compromise and can also result in death. ZIKV can also infect human reproductive tracts leading to sexual transmission. ZIKV infection during pregnancy can cause injury to the placenta and can transmit to the developing fetus, resulting in placental insufficiency, microcephaly, congenital malformations and fetal demise (1), neurological disorders and testis damage (2). Currently, there is no specific treatment for infected patients with ZIKV, but only for the symptoms. Having a better understanding of how ZIKV interacts inside the cell, could lead us to a better design for drugs that prevent or inhibits infection by these viruses, by including protein targets from the host cell. ZIKV is classified taxonomically as a member of the Flaviviridae family that affects mammals and birds (3). ZIKV replication cycle starts when a mature virus enters the cell via membrane receptor, and it is encapsulated into an endosome where undergoes pH acidification by the V-ATPase (4). The endosome breaks and the ZIKV 11 Kb single-strand positive RNA genome is released. The positive RNA translates to a single polyprotein into endoplasmic reticulum (ER) related membranes. The polyprotein then suffers cleavages by either host or viral proteases to produce 10 different proteins; 3 structural proteins: the envelope protein (pE), the membrane protein (prM), and the capsid protein (pC), and 7 non-structural proteins (NS1, NS2A, NS2B, NS3, NS4A, NS4B, and NS5). Mature ZIKV is composed of the 3 structural proteins embedded in the viral membrane: pE, prM, and pC. The +RNA is compacted inside the viral membrane (5). Since viruses in general are big non-motile molecules, they hijack the host cytoskeleton molecular motors in order to move around the eukaryotic cell and complete their replication cycle (6). Human cytoplasmic dynein-1 (dynein) is a 1.5 MDa (1,500 kDa) molecular motor that, together with its cofactor, dynactin, transports different cargoes to the minus end of the microtubules (MT) in an ATP dependent manner (7). Dynein is a homodimer composed of 6 different proteins: the heavy chain (HC), which acts like a scaffold where the intermediate chain (IC) and the light intermediate chain (LIC), bind. The light chain 8 (LC8), light chain 7 (LC7 or roadblock) and the t-complex testis-specific protein 1 (Tctex1) bind to the IC (8). The HC contains a AAA-motif, that hydrolyzes ATP to produce conformational changes in the microtubule-binding domain (MBD) for its binding-unbinding from the MT (9). Dynein-dynactin complex recruits several cargo adaptors to cope with the transport of different cargoes such as single proteins or protein aggregates, endosomes, melanosomes, autophagosomes, lipid droplets, peroxisomes, mitochondria, endoplasmic reticulum, Golgi apparatus, and the vesicles derived from these organelles (6). Dynein also participates in the replication cycle of a great number of viruses from different families including the Flaviviridae family (10). This interaction could be both, by direct interaction with the virus or by interacting with the virus contained in a vesicle. Both cases result in the transport of the viral particles into the perinuclear area over the microtubules (6). There is evidence that shows that ZIKV infection induces massive changes in the host microtubules and intermediate filaments. In this condition, the cytoskeleton produces a microtubule-dependent cage that surrounds the viral factories. Also, by adding paclitaxel, a drug that stabilizes microtubules against depolymerization, the virus does not seem to replicate (11). The flavivirus Dengue virus (DENV) is in close contact with dynein since there is evidence of co-immunoprecipitation and colocalization of the fluorescence labelled anti-DENV E protein and anti-dynein antibodies (11). In this work, we have shown that ZIKV bonds, via its E protein, directly to the dimerization domain of the HC of dynein in a non-vesicle nor dynactin fashion. Also, we have demonstrated the kinetics of this interaction in cell culture indicating that it is exclusively with new-synthetized viral protein. These results strongly suggest that dynein participates in a new cytoplasmic-based step in the current understanding of the ZIKV replication cycle, which is a potential drug target against the infection by the ZIKV.

## Results

### Naturally occurring dynein binds to the ZIKV *in vivo*

#### Envelope-dynein colocalization assay

In Figure 1A we show the ZIKV Envelope protein-dynein colocalization pattern at 8 h, 12 h, and 24 h. The dynein in mock-infected cells was uniformly distributed in the cells, while in the infected cells, clear zones of greater intensity are seen around the nuclei, even in the cells in which the envelope protein is poorly visible, this effect was previously reported by Shrivastava et al 2011. Throughout the experiment, we have shown a notable accumulation of dynein around E protein, which is observed in stoichiometric inequality, favoring dynein in quantity. At 8 h, E protein is shown in low quantity. As time progressed, the amount of E protein increased and at 12 h (Figure 1B), colocalization zones distributed in the cytoplasm were observed with a Pearson’s colocalization index of 0.672. The colocalization distributions starts to accumulate at the perinuclear zone, which is more evident at 24 h. At 24 h (Figure 1C), the Pearson’s colocalization index is 0.698, but the colocalization pattern is localized at the perinuclear zone (Figure C, lower panel), this coincides with the synthesis and assembly of viral proteins near the ER, the distribution of the localization patterns ranging from the cytoplasm to the nucleus as a retrograde traffic, it is necessary to highlight that this is the same pattern observed by Shrivastava et al from 12 to 24 h. Besides, we performed kinetics of dynein and E proteins during the infection (Figure 2) and analyzed both proteins at 8 h, 12 h and 24 h post-infection by WB. At 8 h the envelope protein is undetectable while the dynein is observed with low intensity, but at 12 h the envelope begins to be detectable while the dynein presents a greater intensity in its signal. Finally, at 24 h the envelope protein increases its signal, but the dynein seems to decrease, although the stoichiometric relationship does not seem to be the same, the kinetics seems consistent with what was observed in colocalization, these results are also consistent with our colocalization and the colocalization shown by Shrivastava et al. The results suggest that in an early stage of the infection, dynein would have the task of transporting ZIKV in the by interacting directly with the E protein, for which it would necessarily be overexpressed. However, when this stage is finished, dynein expression goes back to the non-infected levels.

**Fig. 1.**
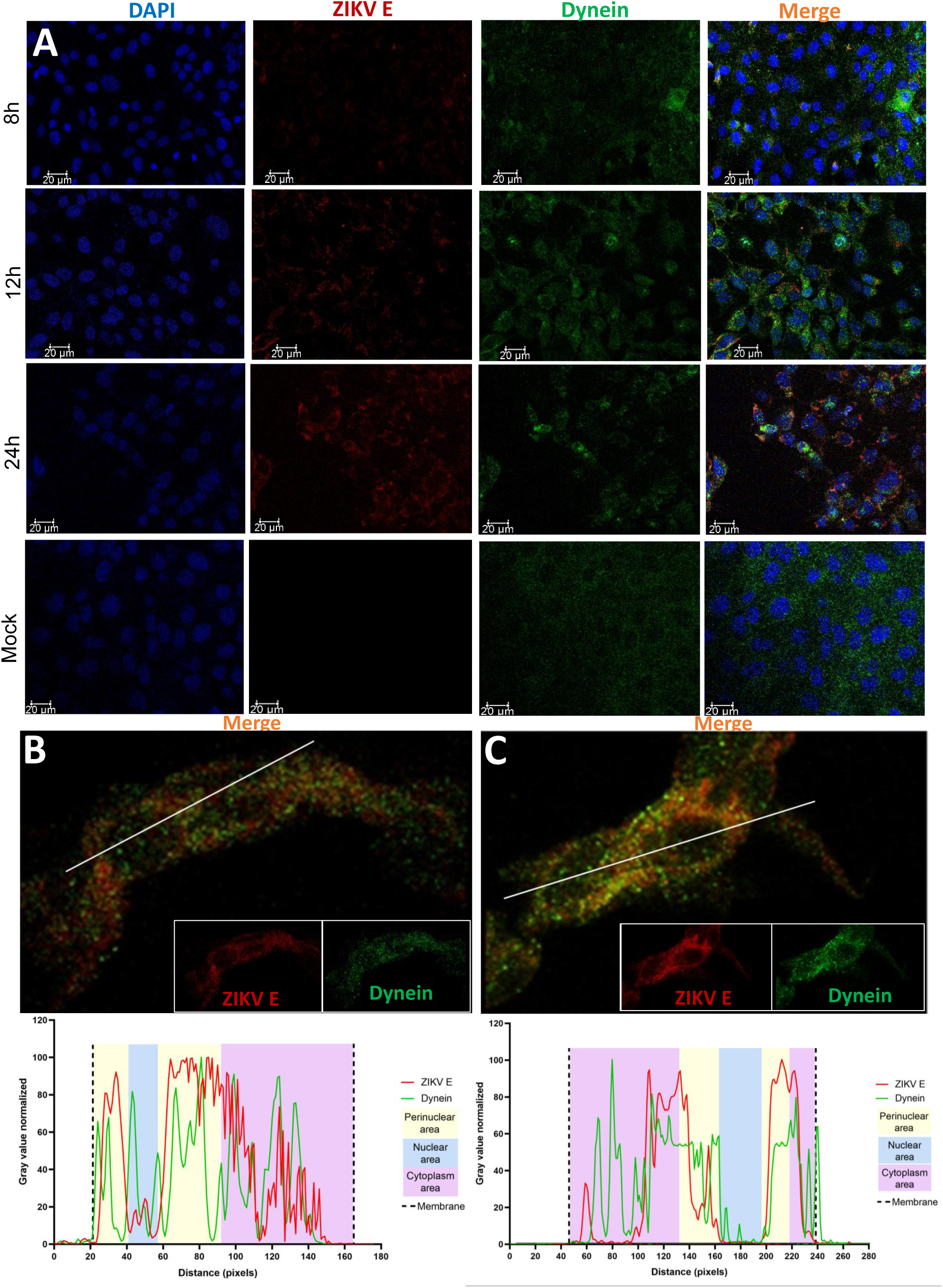
Immunofluorescence Assay of ZIKV infected cells. A. Vero cells were infected with ZIKV and fixed at different times post-infection (p.i.). ZIKV pE was marked with mouse MAb and a secondary antimouse IgG Alexa 488 (red). Dynein was marked with rabbit polyclonal Ab and an anti-rabbit IgG Alexa 594 (green). Nuclei were stained with DAPI (blue). Split images showing the colocalization in a 12, 24, and 48 h time course. Mock cells are cells without infection fixed at 48 h. B. Magnified immunofluorescence image at 12 hours post-infection. C. Magnified immunofluorescence image at 24 hours post-infection.The lower panel of figure B and C shows the colocalization profiles of the sections marked with the white line. The green line is dynein and the red line is pE from the ZIKV. The coloured areas are: Light yellow perinuclear section, blue is the nuclear area, magenta is the cytoplasm area and the dotted line is the plasma membrane

**Fig. 2.**
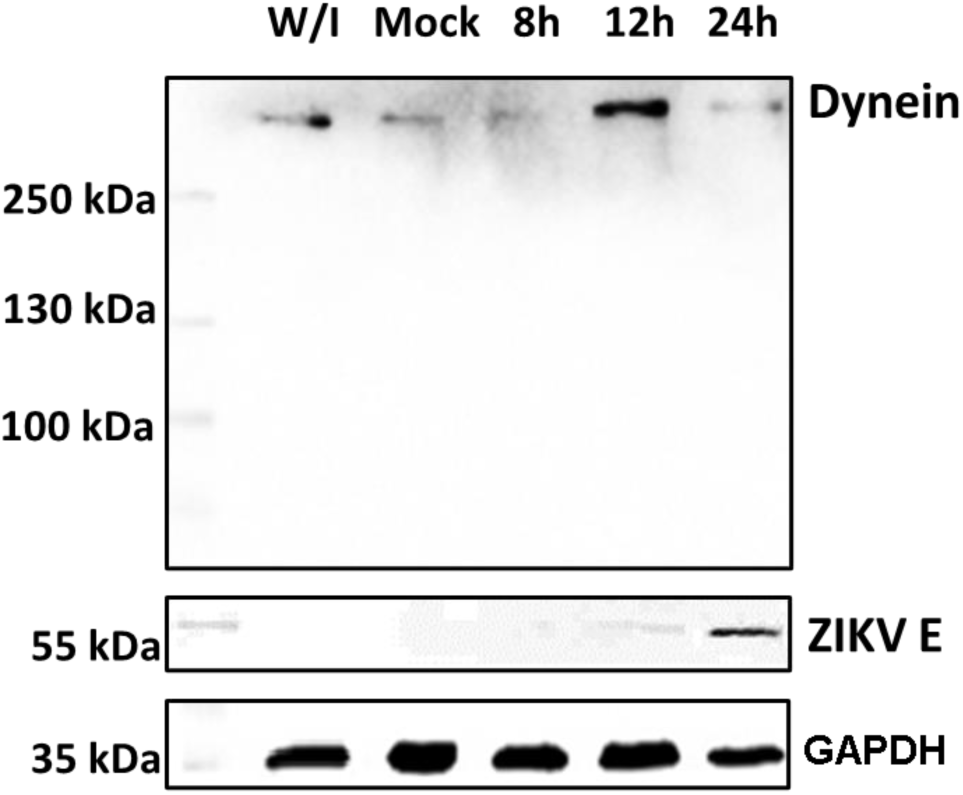
Envelope and dynein kinetics. Line 1. MW marker. Line 2. Vero cells without infection. Line 3.Mock infected, Vero cells infected with inactivated ZIKV. Line 4. Harvested Vero cells 8 h p.i. Line 5. Harvested Vero cells 12 h p.i. Line 6. Harvested 24 h p.i.

#### Envelope-dynein proximity ligation assay

With the proximity ligation assay (PLA), we are evaluating the interaction of proteins at distances <40 nm. ZIKV infected cells were fixed at 12, 18, 24 and 48 hpi. (Figure 3). Weak positive signals (1 or fewer PLA dots per cell) for dynein E protein interactions were observed at 12 and 24 h p.i. but clear positive signals (6 or more dots per cell) were observed in cells fixed at 18 hpi. The absence of any signal in the 2 negative controls used corroborated the specificity of the assay. These results agree with the colocalization and protein expression results, which suggest that in ZIKV infected cells, dynein and the E protection interact in a narrow time window. The loss of the interaction after 18 h, could mean that the ZIKV is being processed inside vesicles, so there is no need for a direct interaction of dynein with the virus.

**Fig. 3.**
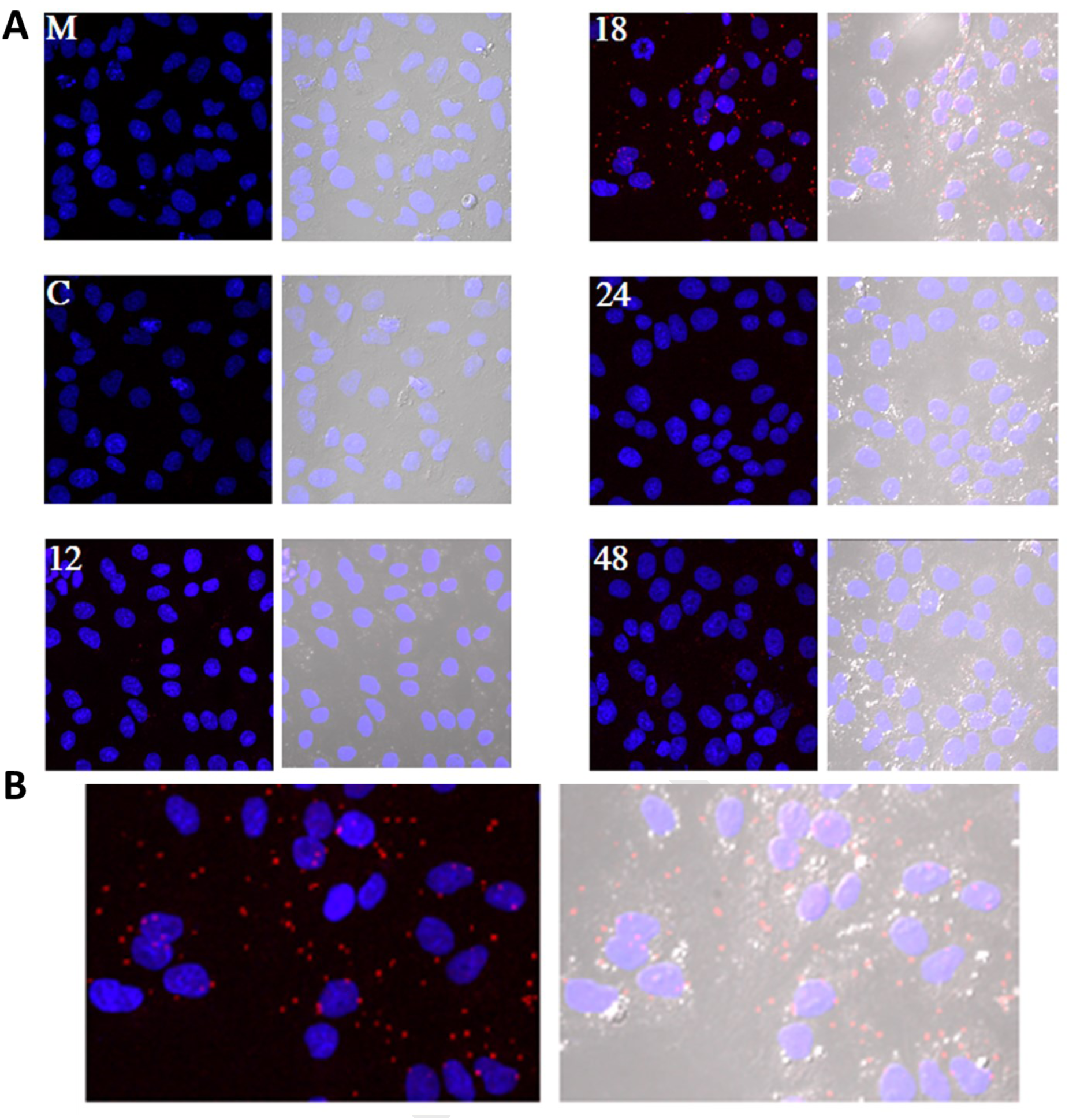
Proximity ligation assay of ZIKV infected cells. A. (M) Mock image is cells without infection fixed at 48 h (C) control was cells infected with the virus at 18 h without the incubation of the anti-pE antibody. For the time course, we used Vero cell infected with ZIKV and fixed at different times post-infection: 12, 18, 24, and 48 h This assay was performed with a monoclonal anti-pE in mouse MAb and with a polyclonal anti-NTDyn in rabbit. The PLA dots were marked with Cy3 (red). The images on the right show DAPI (blue) and Cy3 (red) channels. The images on the left show blue, red and the differential interference contrast channel (grey). B. Magnified image at 18 h post-infection.

#### Zika virions interact with NTDyn

In order to examine whether the complete ZIKV is interacting with the NTDyn recombinant protein, we bound the His6NTDyn into a His Trap Nickel column and then we injected an enriched sample of ZIKV, harvested from infected Vero cells. We washed the column, eluted the bound material with imidazole and analyzed the fractions by SDS-PAGE (Figure 5A); lane 1MW marker, line 2 the purified His6NTDyn, in lane 3 the purified ZIKV, lane 4 eluted material where we can see the band corresponding to His6NTDyn and some bands corresponding to the MW of the E protein. With the idea of delimiting the interaction region, we injected the recombinant His6NDD (Figure 5B), this is the central core of the His6NTDyn fragment (Figure 6B). In line 1 MWM, line 2 is the purified His6NDD, line 3 purified ZIKV and line 4 elution shows the presence of both bands, this result is consistent with the previous result, and it delimits the zone of interaction between ZIKV and dynein, this is in the dimerization domain. As a negative control, we used a polyclonal anti-NDD antibody to block this domain in the His6NTDyn fragment, to expose only the helical bundles 1, 2 and 3 of dynein’s HC, this complex was injected into an affinity column and subsequently washed and injected with the ZIKV particle, and eluted. In Figure 5C we observe in line 1 MW marker, line 2 NTDyn and anti-NDD antibody (IgG) bands, line 3 band corresponding to the ZIKV envelope protein in line 4 elution where we only observe the bands corresponding to the His6NTDyn-Anti NDD complex, the absence of bands corresponding to the envelope, shows the absence of ZIKA-complex interaction, this result shows the specificity of the ZIKV-NDD interaction. As a control, we injected our purified Zika viral particle into an affinity column, to analyze the capacity to form unspecific interactions (Figure 5D). We also observed in line 1 MW marker, line 2 envelope zika protein band, and line 3 elution band, where no band is observed, this experiment confirms that ZIKV does not interact non-specifically with the affinity column.

**Fig. 4.**
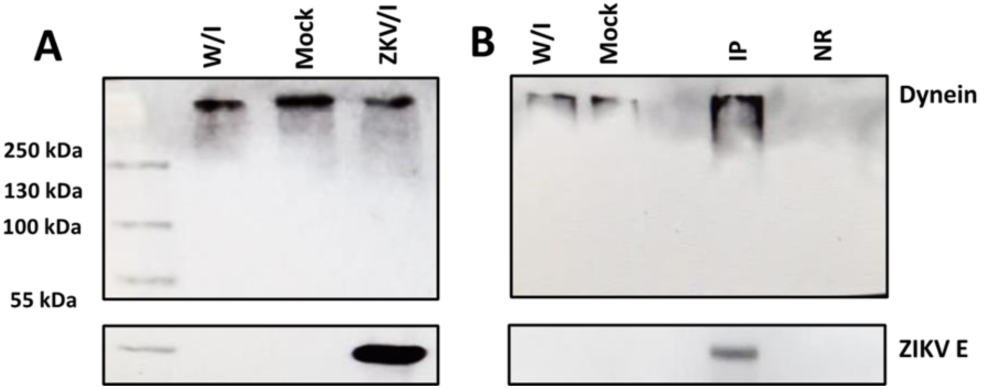
Binding of the ZIKV with NTDyn. A Purified ZIKV from Vero cells was loaded into a 10 ml HisTrap HP Nickel (GE) column pre-loaded with the His6NTDyn. The column was washed, and the sample was eluted with imidazole. The fractions were analyzed with an SDS-PAGE. Line 1. MW marker. Line 2. The purified His6NTDyn band. Line 3. Enriched pE from the ZIKV. Line 4. The eluted fraction contains both, the complete ZIKV and NTDyn-His6. B. Line 1. MW marker. Line 2. His6NDD purified protein bound to affinity column, Line 3. Purified ZIKV. Line 4. Elution, C. Line 1. MW marker. Line 2. His6NTDyn + anti-NDD antibody complex. Line 3. Purified ZIKV. Line 4. Elution. D. Line 1. MW marker. Line 2. Purified ZIKV. Line 3. Elution.

**Fig. 5.**
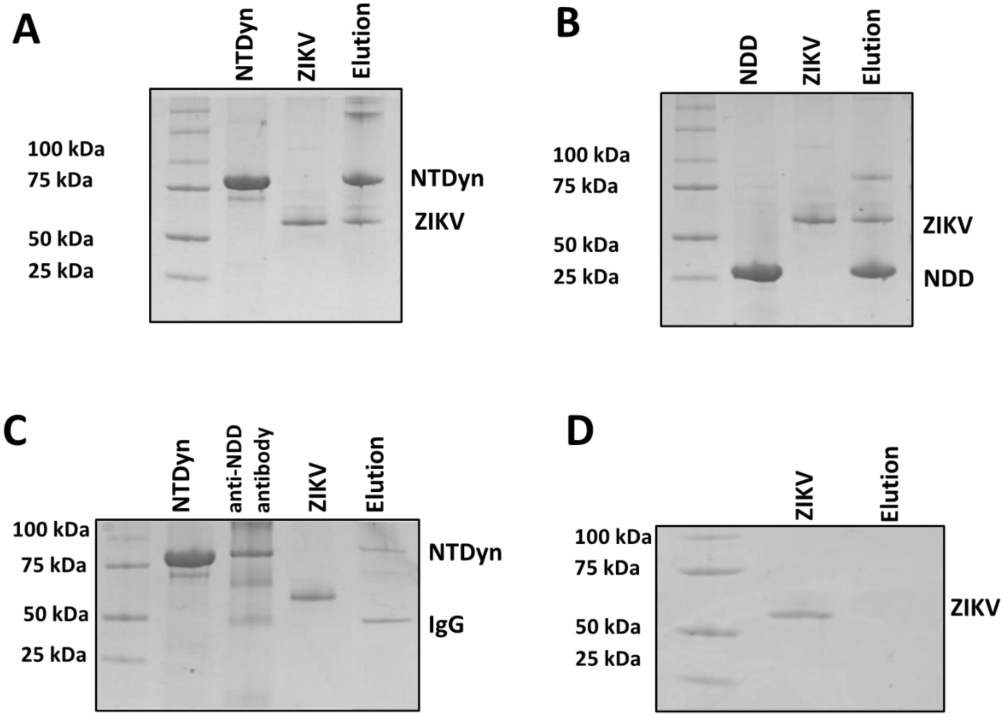
Binding of the ZIKV with NTDyn. Purified ZIKV from Vero cells was loaded into a 10 ml HisTrap HP Nickel (GE) column pre-loaded with the His6NTDyn. The column was washed, and the sample3 was eluted with imidazole. The fractions were analyzed with an SDS-PAGE. Line 1. MW marker. Line 2. The purified His6NTDyn band. Line 3. Enriched pE from the ZIKV. Line 4. The eluted fraction contains both, the complete ZIKV and NTDyn-His. Line 1. MW marker. Line 2. His6NDD purified protein bound to affinity column, Line 3. Purified ZIKV. Line 4. Elution, C. Line 1. MW marker. Line 2. His6NTDyn + anti-NDD antibody complex. Line 3. Purified ZIKV. Line 4. Elution. D. Line 1. MW marker. Line 2. Purified ZIKV. Line 3. Elution

**Fig. 6.**
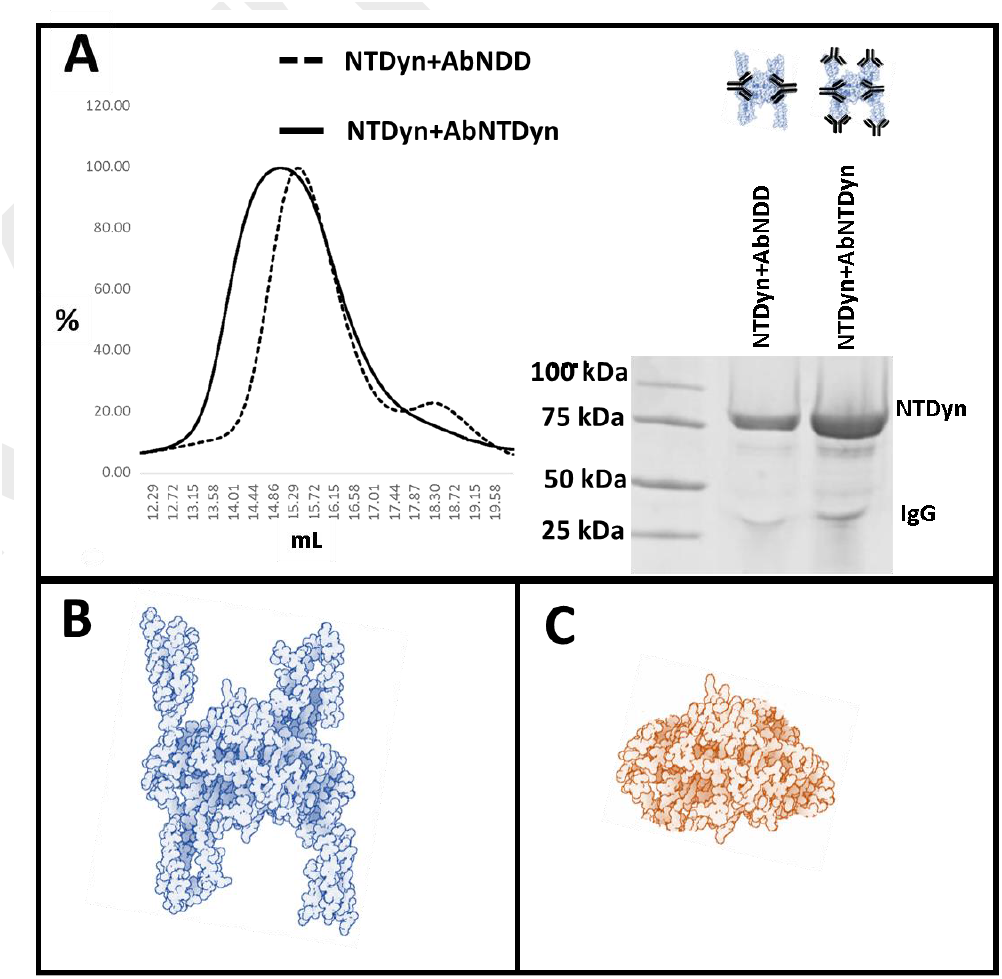
NTDyn-antibodies complex A. size-exclusion chromatogram, of the NTDyn-anti-NTDyn and NTDyn-anti-NDD complexes on the right side the SDS-PAGE of the corresponding elution peaks, is shown. B. left panel, the structure of the blue NTDyn is shown, composed in its nucleus by the NDD and in its external part by 6 lateral alpha-helices. Modified images of 5NW4(21) and 5OWO(22). C. NDD. Dimerization domine of dynein in red color. Modified images of 5OWO.

#### NTDyn antibodies complex size-exclusion chromatography profiles

To verify that the anti-NDD antibody was binding only to the NDD of the NTDyn fragment, we analyzed this complex by gel filtration and compared it with the NTDyn-anti-NTDyn complex. In this way, we were able to analyze the interaction capacity of the antibody since both polyclonal antibodies have different binding sites hence the NTDyn-anti-NTDyn complex would show a reduction in its elution volume compared to the NTDyn-anti-NDD complex. Because the NTDyn-anti-NDD complex -NDD would not have anti-bodies bound to the alpha-helices of the NT-Dyn fragment. Which would give us the ability to test the interaction between the alpha-helices of NTDyn and ZIKV. As a result, we observe in figure 6A, right panel, the chromatogram corresponding to the Size-exclusion chromatography of both complexes, where it is shown that the maximum elution peak of the NTDyn-anti-NDD complex occurred at 15.02 ml. While the maximum peak of the NTDyn-anti-NTDyn complex occurred at 15.39 ml, this result is consistent with expected, since the NTDyn-anti-NDD complex should have fewer bound antibodies, so it would show a lower elution volume. The chromatography fractions were analyzed by SDS-PAGE, where the presence of bands corresponding to the NT-Dyn fragment and IgG was observed.

## Discussion

The recent SARS CoV-2 outbreak has been an example of how fast viral infections can be globally spread in a minimal fraction of time, and how extremely vulnerable to the action of the virus we are. The virus has a high mutation rate, this helps the virus to cope with antivirals and to gain resistance against them. Finding host cells’ molecules that interact with the virus and targeting that interaction with new drugs could diminish the probability of antiviral resistance. In this work, we evaluate the role of human cytoplasmic dynein-1 (dynein) in the replication cycle of the Zika virus (ZIKV) and obtained clear evidence that indicates that those two proteins interact in vitro as well as in infected cells. First, we investigated whether this interaction could be present in infected cells, by infecting Vero cells, fixing samples at 12, 24, and 48 h post-infection. We revised the immunolocalization of the ZIKV and the naturally occurring dynein, the maximum colocalization was at 24 hours, this is consistent with the reported initiation of viral protein synthesis in this cell line (12). The colocalization of fluorophores in confocal microscopy could have far interactions 400-600 nm due to the limited resolution of the optical system. To increase the resolution of our method, we used the proximity ligation assay, which has a theoretical maximum limit of 40 nm. There are some limitations or false interpretations of this technique (13). On the other hand, during the transport of the virus towards the perinuclear zone, the virus is transported in the endosome. For this, the necessary machinery is GTPase Rab 5 or 7, which regulates transport to early or late endosomes, respectively (14). These membrane proteins of the endosome interact with the cargo adaptor on one end, and on the other end, it interacts with the dynein-dynactin complex (15). The endosome membrane proteins, and the coiled-coil cargo adaptor whose length we have calculated to be approximately 66 nm (16), would not allow us to obtain a signal in the PLA (<40 nm). In addition, during immunoprecipitation, the RIPA buffer solubilizes the lipids from the native membranes, since ZIKV is inside the vesicle, in case the interaction that we are capturing could be with the ZIKV inside a membrane vesicle, the virus would be released of the vesicle, and it will not interact with dynein. Since we are still detecting dynein-ZIKV interaction with the immunoprecipitation experiments, it means that we are capturing a complex with direct interaction. We have also performed kinetics of the infection, with a maximum of PLA signal at 18 h p.i. and after this time, we observed a decrease of the PLA dots. We increased the MOI from 1 to 5 in order to check some insights of direct interaction at the beginning of the replication cycle with no positive PLA at 12 h or less, nor after 24 h (data not shown). We propose that after 18 h, the viruses are being processed inside vesicles, so the direct interaction between ZIKV and dynein is not detectable. Taking together, our data led us to propose the next addition on the ZIKV replication cycle; since the ZIKV release of the first new virions is approximately 24 hours post-infection, and we are losing both, colocalization and PLA signals after 18 hours, we believe that ZIKV’s replication cycle step on which dynein participates should be when newly synthesized viral proteins in the cytoplasm (11). The co-localization assay and PLA could only determine closeness and no interaction in vitro, which led us to analyze the interaction through the immunoprecipitation of the complex, this assay in infected Vero cells guarantees that this interaction occurs naturally during the viral replication cycle. We propose three possible scenarios; first, we suggest that the ZIKV polyprotein starts translating on the cytosol, where it hijacks the infection-dependent highly expressed dynein (12-18 h) that transports the full or partially non-processed polyprotein to the viral factories (not direct dynein-ZIKV interaction), where the virions will be processed, assembled and then, transported into the Golgi apparatus to its final maturation step before it is released to the extracellular space (24 h). There is evidence in the literature of a non-processed polyprotein composed of C-prM-E-NS1 proteins of the YFV flavivirus, synthesized with the in vitro translation system in rabbit reticulocyte lysates (17). This translation system in rabbit reticulocyte lysates has no microsomal membranes. The second scenario is that the dynein-ZIKV complex could be formed due to the translocation of factors from the ER lumen to the cell surface that could facilitate in yet unknown ways apoptotic signalling cascade (8). It has been observed that, under stress conditions, the permeability of the ER allows luminal proteins to be released or translocate to the cytoplasmic side of the ER (18). This would also require the transport of E protein by the dynein to the viral replication factories. The third and final scenario we are proposing is a dual polyprotein topology an unknown process of post-translational translocation leading to the non-uniform topology, where there is an equilibrium of envelope protein copies in luminal or in cytosolic compartments (8). In hepatitis B virus (HBV) has been observed that all envelope proteins synthesized in transfected cells or in a cell-free system adopt more than one transmembrane orientation (**?**). In this way, vesicles with E protein from ZIKV facing cytosol could bind to dynein and the whole vesicle will be transported by the molecular motor in order to reach the viral factories. Thus, we have shown strong evidence of the first non-coiled-coil protein that interacts with dynein without a cargo adaptor or dynactin. In order to prove this strong interaction with the complete Zika virion, we attached the His6NTDyn into an affinity Nickel column and then, we bound an enriched extract of virions from infected Vero cells to this column and elute the samples. In Figure 4A we show that NTDyn was able to bind Zika virions since both molecules co-elute with imidazole. Once we eliminate the helical-bundles 1, 2 and 3 (residues 202-504) of the NTDyn by using the NDD, we carry out the same experiment where we observe the same result the obtained with the NTDyn, this delimits the interaction to the first 201 amino acids of the amino-terminal fraction of the heavy chain of dynein. This could be the answer to why the ZIKV binds to dynein without dynactin or cargo adaptor; dynactin and the cargo adaptors BICDR, BICD2 and HOOK3 bind to the helical bundles 1, 2 and 3, the interaction we are proposing is in the ‘opposite face’ of dynein. Although we do not know the mechanism of this interaction and whether ZIKV binding promotes dynein processivity, we suggest that it is a retrograde transport function. In addition, the position of NDD in the complex would give it the ability to interact with the ZIKV envelope, perhaps during its retrograde movement (Figure 7). In comparison with similar studies, such as Brault et al, where they characterize an interaction between Tctex-1 and prM, which could be attributed to a different biological role, due to the ability of Tctex-1 to lodge in different cellular compartments (19). Shrivastava et al, attribute a retrograde trafficking role to an observed interaction between the LC8 dynein light chain and DENV pE, although does not clearly address the location of the interaction (20).

**Fig. 7.**
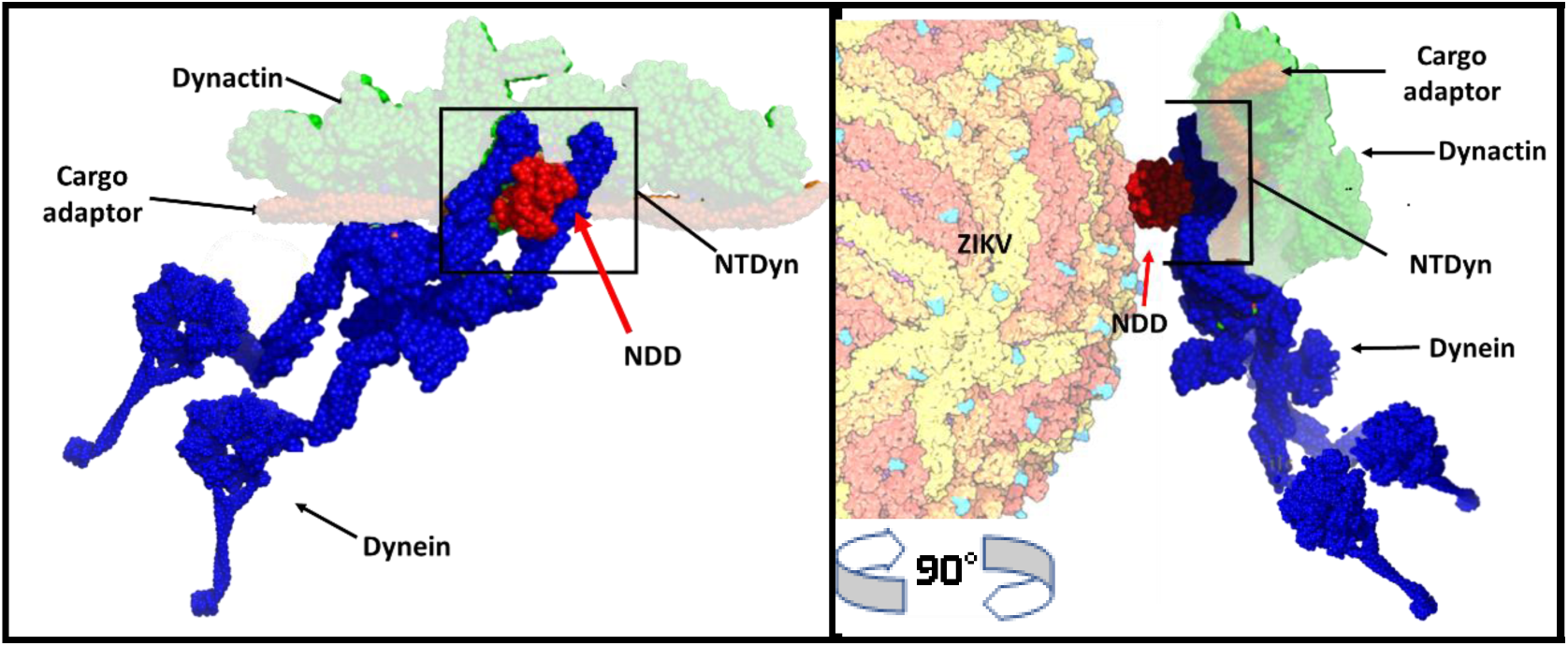
NDD is exposed to ZIKV interaction in a retrograde traffic complex. In blue, dynein forms a complex with dynactin in green, and in brown, the cargo adapter. We emphasize the position of the NDD in red within the complex and its ability to be recruited by the envelope protein. PDBs 3VKH (23) and 5NW4(21).

## Conclusions

We have shown strong experimental evidence that demonstrates the interaction between the E protein from the Zika virus and the dimerization domain of human dynein, this interaction occurs during the viral replication cycle between 18 and 24 h post-infection, this interaction is an uncharacterized step in the viral replication cycle that could be targeted to design a new drug that stops virus spreading into the currently infected patients.

## ACKNOWLEDGEMENTS

The authors of this paper want to thank Dr. Andrew Carter from the LMB-MRC, Cambridge, UK, for the generous gift of NDD and Nt-Dyn bacterial expression vectors; Prof. Glaucius Oliva from the University of Sao Paulo, Brazil, for the generous gift of the complete ZIKV envelope protein expression vector; Saul Santiago Sánchez for their help with the production of polyclonal antibodies; Enrique Ahmed Alcaraz Perez for his valuable technical support throughout this investigation; Henriques, R. for the Biorxiv template. EMR acknowledges funding from Instituto de Bebidas para el Bienestar y la Salud (Premio de Investigación en Biomedicina “Ruben Lisker” 2017), CONACYT (CB2017-2018: A1-S-10743), SEP-CINVESTAV (2018:1), Agencia Mexicana de Cooperación Internacional para el Desarrollo (AMEXCID) of the Secretaría de Relaciones Exteriores (SRE) (Project AMEXCID 2020-3) and Gobierno del Estado de Hidalgo (Proyecto Sincrotrón 20201120). DIZV, GVC, RRR and JGFV received a pre-doctoral fellowship from CONACYT.

## Materials and Methods

### Immunofluorescence Assay

Vero cells were seeded one day prior to infection at a minimum concentration of 1×10^5^ cells/ml in cell culture flasks. Cells were infected with ZIKV (MR766 strain) at a multiplicity of infection (MOI) of 1. Adsorption was carried out for 1 h at 4 °C to prevent virus entry and synchronize infection of cells. Cells were washed twice with a cold medium to remove unbound virus and replenished with a prewarmed medium. The coverslips were taken out of the culture at 12, 48 and 72 h postinfection (p.i.) under sterile conditions, washed with 0.01 M phosphate buffer saline pH 7.2 (PBS) and fixed in ice-cold acetone for 10 min at 20 °C for experiments. The fixed cells were washed with PBS and blocked with 1% bovine serum albumin (BSA) in PBS for 30 min. For dual staining, NTDyn was labelled with our lab-made polyclonal antibody (rabbit) and the E protein using the pan-flavivirus monoclonal antibody 4G2. The primary antibodies were added simultaneously and incubated for 1 h followed by washing with PBS. This was followed by incubation with appropriate secondary antibodies conjugated to fluorescent dyes, Alexa 488 or Alexa 594 added simultaneously. DAPI (4, 6 diamino-2-phenylindole) was used to stain the nucleus in all experiments. At the end of the staining process, coverslips were washed and mounted onto slides using a mounting medium. Control samples were non-infected cells processed with the same procedures described previously and were included in all experiments as mock-infected cultures. Cells were observed and images were digitalized in a Zeiss 700 Confocal microscope.

### Envelope and dynein protein expression kinetics

Vero cells at 80% confluency were infected with ZIKV at an MOI of 5. Infection kinetics at 8, 12 and 24 hours were performed. Uninfected Vero cells and Vero cells treated with a heat-inactivated virus (mock) were used as controls. Cells were then lysed with RIPA buffer (100 mM Tris-HCl pH 8.3, 2% Triton X-100, 150 mM NaCl, 0.6 M KCl, 5 mM EDTA) in the presence of one tablet of the protease inhibitor cocktail per 25 ml (Complete, Invitrogen). Cell lysates were analyzed by 12% SDS-PAGE for 80 min at 100 V (Mini-Protean Cell, Amersham Biosciences, Piscataway, NJ, USA) and transferred into nitrocellulose membranes (BIO-RAD) was carried out at 120 V for approximately 2 hours). Membranes were blocked with PBS-Tween-5% milk for one hour and then washed four times with PBS-Tween. Membranes were then incubated overnight with the primary antibodies at 4 °C. After this incubation, the membranes were washed again with 1X PBS and incubated with HRP-coupled secondary (Invitrogen) for one hour. After another round of 4 washes with PBS-Tween, the membranes were developed in the presence of the chemiluminescence developer reagent (Thermo Scientific) following the manufacturer’s instructions, using the ChemiDoc kit (BIO-RAD) and digitalized with the Chemi-Doc XRS System (BIO-RAD).

### Proximity ligation assays

Vero cells were seeded one day prior to infection at a minimum concentration of 1×10^5^ cells/ml on glass coverslips in a 24 well plate. Cells were infected with ZIKV (MR766 strain) at a MOI=1. Then cells were fixed with formaldehyde 4% for 10 min at 8, 12, 18, 24 and 48 h p.i. After this, the cell culture was washed with wash buffer (PBS Tween 0.1%) and permeated with PBS-Tween 0.2% for 10 min. Once removed the permeabilization buffer, samples were blocked with blocking buffer (5% bovine serum albumin in PBS) for 1 hour at 37 °C and continued with wash and incubation with the primary antibodies 1:100 dilution overnight at 4° C with blocking buffer. Mock-infected cells were incubated with both primary antibodies and control infected cells were incubated without the anti-E mAb, both samples were run in parallel as negative controls. The next day, samples were washed, and the PLA probe was added in 1:5 dilution with blocking buffer for 1 hour at 37°C in a humidity chamber. The sample was washed and incubated with the ligation mix (6 µL of the concentrated ligation buffer with water for a total reaction volume of 30 µL and 1 µL of ligase) for 30 min at 37 °C. Then, the sample was washed and incubated with the amplification mix (6 µL of the concentrated amplification buffer with water for a total reaction volume of 30 µL and 0.5 µL of polymerase) for 100 min at 37 °C. After this incubation, the cells were washed with wash buffer B and the sample was then incubated with 4,6-diamidino-2-phenylindole (DAPI) to stain nuclei for 10 min under gentle agitation. Next, the samples were washed one time with B buffer 1X and one time with buffer B 0.01X for 1 min. Finally, the samples were mounted and analyzed in a confocal microscope (Zeiss 700). Results were quantified by ImageJ and expressed as several dots/cells.

### Immunoprecipitation envelope protein–dynein

Vero cells were seeded one day prior to infection in cell culture flasks. The cells were infected with the ZIKV MR766 strain at a MOI of 5. The medium was removed 24 h later and cross-linked with 1 70% formaldehyde and incubated for 10 minutes at 37 ° C, cross-linker was removed washing twice with cold PBS 1X. Cells were incubated with RIPA buffer for one hour on ice, then the sample was centrifuged at 13000 g for 20 min at 4 °C and the supernatant was recovered. 20 µL of magnetic beads were taken and washed four times with solution A (25 mM Tris HCL pH 7.5, 125 mM NaCl, 2.5 mM EDTA, 2.5 mM EGTA, 2.5 mM NaF, 0.1 70% Triton X100). 10 µL of Antibody (Anti-E, 4G2) were added and incubated overnight at 4 °C. The next day, the supernatant of the samples was added, and the sample was incubated overnight at 4 °C. The magnetic beads were then retrieved using the magnet and washed four times with wash solution B (25 mM Tris HCL pH 7.5, 125 mM NaCl, 2.5 mM EDTA, 2.5 mM EGTA, 2.5 mM NaF, 0.5% Triton X100). Finally, 50 µL of sol C (25 mM Tris pH 8.0, 2.5 mM EDTA, 0.1% SDS) are added and incubated at 65 °C for 30 minutes to recover the supernatant, which was analyzed by Western Blot with the Mab Anti-E, Pab Anti-NTDyn.

### ZIKV envelope protein expression and purification

The complete *E. coli* codon-optimized coding sequence for the ZIKV envelope protein (E) (corresponding to UVE/ZIKV/1947/UG/MR766) was a generous gift from Prof. Glaucius Oliva from the University of Sao Paulo. The DNA corresponding to the E-protein without the membrane domain (Ep) was subcloned to pRSET A plasmid with an N-terminal His6-tag. This version of E protein was expressed in *E. coli* SoluBL21 strain (Genlantis) into inclusion bodies (IB). The E protein was purified as in (Dai et al, 2016) with the following modifications, we started the solubilization of the IB with 8 M urea 6 h at room temperature. Subsequently, the solubilized E protein was applied into a Histrap HP 5 ml pre-equilibrated with the refolding buffer (100 mM Tris pH 8.0, 400 mM L-Arg HCl, 2 mM EDTA, 5 mM reduced glutathione, 0.5 mM oxidized glutathione, and 5% glycerol) and then eluted with the refolding buffer with 500 mM imidazole. The His6-tag was cleaved with the TEV protease and concentrated by centrifugation (1650 g at 4 ºC) in an Amicon Ultra 15 concentrator with 30 kDa cutoff. 500 µl of the concentrated E protein, was purified further with a Superdex 200 10/300 Increase gel filtration column (GE Healthcare). Fractions containing the E protein were pooled and concentrated at 10 mg/ml.

### Expression and purification of the N-term Dynein Heavy chain and dimerization domain

Both proteins were individually purified using the same protocol. The *E. coli* codon-optimized human dynein heavy chain N-Terminus residues 1-560 (NTDyn) and human cytoplasmic dynein dimerization domain 1-201 (NDD) with N-terminal His6-tag and TEV protease cleavage site into the pRSET (A) plasmid was expressed in *E. coli* SoluBL21 strain as described previously with some modifications (Urnavicius, et al. 2018). Once expressed, the cells were sonicated (500W Ultrasonic processor Cold Palmer CV334), with 5-35 seconds pulse-rest cycles for 3 minutes at 50% power with buffer A (30 mM Tris pH 7.5, 200 mM NaCl, 2 mM MgCl_2_, 25 mM imidazole, 1 mM benzamidine, 1 mM PMSF, 1 mM beta-mercaptoethanol). Next, the cell lysate was ultra-centrifugated at 165,000 xg for 30 minutes at 4 °C; the clarified material was filtered with a 0.22 µm membrane and then applied a HisTrap HP 5 ml (GE Healthcare) and eluted with buffer B (30 mM Tris pH 7.5, 200 mM NaCl, 2 mM MgCl_2_, 500 mM imidazole, 1 mM benzamidine, 1 mM PMSF, 1mM beta-mercaptoethanol). The eluted protein was desalted with a desalting column (HiPrep 26/10 GE) and digested for 18 h at 4 ºC with the TEV protease. The removed His6-tag and the TEV protease were bound to the HisTrap affinity column, and the protein was eluted in the washing step. The fractions were concentrated to 500 µl by Amicon Ultra 15 concentrators with 30 kDa cutoff and applied to a Superdex 200 10/300 Increase gel filtration column (GE Healthcare). Individual fractions containing the NTDyn and NDD were pooled and concentrated at 10 mg/ml.

### NTDyn or NDD-ZIKV interaction on IMAC

For experiments A and B 100 µg of protein His6-NTDyn or His6-NDD was bound to 1 ml HisTrap HP previously equilibrated with phosphate-buffered saline (PBS1X) pH 7.4 and then washed with 5 ml of the same buffer to later bound the viral particle in Molar ratio 10:1 protein: virus. Viral particles were harvested 24 h p.i., purified and injected through the column and subsequently eluted with 5 ml of elution buffer (30 mM Tris pH 7.5, 200 mM NaCl, 2 mM MgCl_2_, 500 mM imidazole, 1 mM benzamidine, 1 mM PMSF, 1 mM beta-mercaptoethanol). For the protein:antibody complexes, 100 µg of NTDyn or NDD interacted with anti-NTDyn or anti-NDD polyclonal rabbit antibodies for 30 min in a 1:4 protein: antibody ratio. Once the complexes were formed, they were bound to the affinity column previously equilibrated with (PBS) pH 7.4 and washed with 5 ml of the same buffer to later bound the viral particle in Molar ratio 10:1 protein: virus. Viral particles were harvested 24 h p.i., purified and injected through the column and subsequently eluted with 5 ml of elution buffer (30 mM Tris pH 7.5, 200 mM NaCl, 2 mM MgCl_2_, 500 mM imidazole, 1 mM benzamidine, 1 mM PMSF, 1 mM beta-mercaptoethanol). After elution, fractions were analyzed by SDS-PAGE.

### NTDyn-antibodies complexes size-exclusion chromatography profiles

Two different experiments were performed, the first was the injection of the NTDyn and anti-NTDyn polyclonal antibody complex, this was injected after a 30 mins incubation in a 1:4 molar ratio NTDyn:antibody to allow interaction. The second experiment consisted of injecting the NTDyn and anti-NDD antibody complex. These experiments were designed to block the NTDyn against the interaction with the ZIKV, using the Anti-NTDyn pAb, and by using the NDD pAb, we are blocking only the NDD but the helical bundles 1, 2 and 3 are still free for the interaction. Experiments were performed using a Superdex 200 10/300 size exclusion column (GE Healthcare) previously equilibrated with PBS 1X pH 7.4 buffer at room temperature at a flow rate of 0.5 ml/min. The resulting fractions were analyzed with respect to output volume and by SDS-PAGE.

## Supplementary material

### A. Anti-NTDyn and anti-NDD pAbs production

Both antibodies were individually produced and used the same protocol. Four 10-12 weeks-old Rabbit NZB Crl:KBL(NZW)BR were obtained from the breeding facilities at the CINVESTAV (Centro de Investigation y de Estudios Avanzados del Instituto Politecnico Nacional). All animals were housed and handled in accordance with institutional guidelines. The rabbits were immunized with three doses of 100 µg of NTDyn or NDD recombinant proteins. These immunizations were administered subcutaneously with complete Freund’s adjuvant at the first inoculation, and with the incomplete Freund’s adjuvant, at the first, third and fifth week. In the seventh week, the rabbits were sacrificed and the antibodies in the serum were purified by protein A-Sepharose 4B (Invitrogen).

### B. Far-Western blot NTDyn-E protein

To address the interaction between the recombinant E protein and NTDyn we did a far-western blot. Initially, we transferred in five lanes of a PVDF membrane 100 µg of the recombinant E protein in each lane. After membrane blockade we made it interact with different amounts of the recombinant NTDyn. As a result, we observed in figure S1 an increase in the intensity of the signal that corresponds to the increase in the concentration of NTDyn. As a control we used a lane transferred with 100 µg of NTDyn to which we did not interact with anything, this allowed us to observe the intensity of the signal generated by the reactivity of the antibody, for the negative control we took a lane transferred with recombinant E protein and made it interact with 100 µg of BSA. As a result, we do not observe a signal, which shows us the specificity of the interaction. These results corroborated our hypothesis and led to the analysis of this interaction between recombinant proteins. Methodology. Five lines of the recombinant purified E protein and one of NTDyn were analyzed by 12 percent SDS-PAGE for 80 min at 100 V (Mini-Protean Cell Amersham Biosciences, NJ, USA) and transferred into PVDF membranes (BIO-RAD) was carried out at 120 V for approximately 2 hours). Membranes were blocked with PBS-Tween-5 percent milk for one hour and then washed four times with PBS-Tween. Membranes were then incubated two hours with 0, 10, 100, 1000 g 120 min at TA with PBS. Then the membranes were incubated overnight with the primary antibodies at 4 °C. After this incubation, the membranes were washed again with 1X PBS and incubated with HRP-coupled secondary (Invitrogen) for one hour. After another round of 4 washes with PBS-Tween, the membranes were developed in the presence of the chemiluminescence developer reagent (Amersham ECL Western Blotting Detection Reagent Cytiva) and digitalized with Amersham Imager 600 System GE Healthcare.

**Fig. 8.**
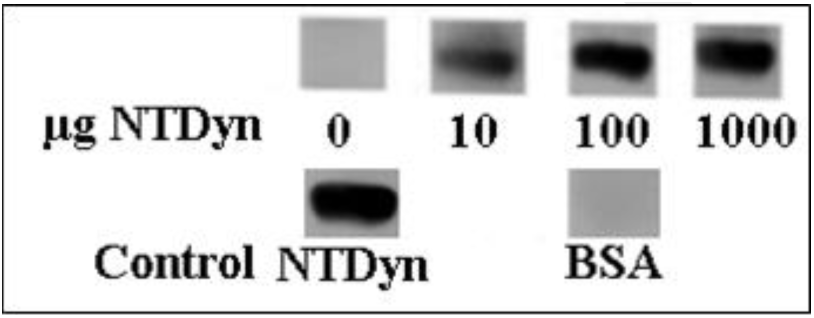
Far western Blot E protein-NTDyn. In the four upper lanes the interaction of the E protein with different concentrations of NTDyn is shown, in two lower lanes the controls of NTDyn and interaction with BSA are shown.

### C. NTDyn and NDD gel filtration chromatography

The NTDyn fragment and the NDD were both purified following the protocol referred in material and methods. Using an affinity chromatography step and a gel filtration step. Finally, the fractions resulting from gel filtration were concentrated and analyzed with SDS-PAGE.

**Fig. 9.**
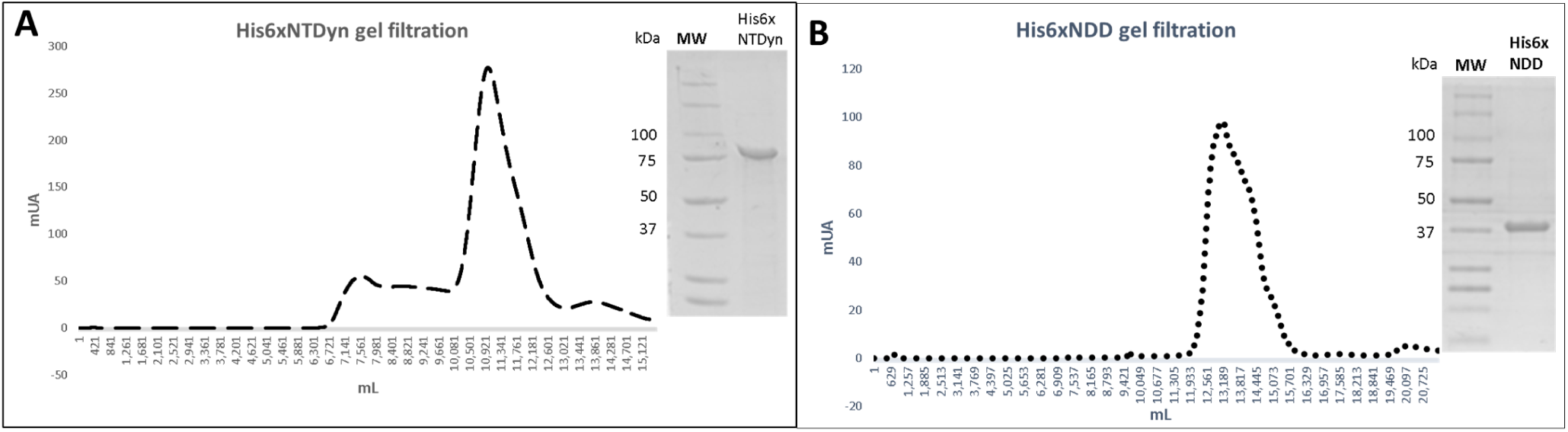
NTDyn and NDD purification. A. NTDyn gel filtration chromatography and SDS PAGE, B. NDD gel filtration chromatography and SDS PAGE

